# Dynein-mediated trafficking and degradation of nephrin in diabetic podocytopathy

**DOI:** 10.1101/2022.10.01.510475

**Authors:** Hua Sun, Jillian Weidner, Chantal Allamargot, Robert Piper, Jason Misurac, Carla Nester

**Author notes:** **Corresponding author:** Hua Sun, Address: 200 Hawkins Drive, Iowa City, IA 52242.

## Abstract

Diabetic nephropathy (DN) is characterized by increased endocytosis and degradation of nephrin, a protein that comprises the molecular sieve of the glomerular filtration barrier, but the key trafficking mechanism that connects the initial endocytic events and the homeostasis of nephrin is unknown. Our work implicates cytoplasmic dynein, a transport complex that is upregulated in DN, plays a critical role in triaging the endocytosed nephrin between recycling and proteolytic pathways. Using *Nephroseq* platform, our transcription analysis in public DN databases revealed dynein overexpression in human DN and diabetic mouse kidney, correlated with the severity of hyperglycemia and nephropathy. The increased expression of dynein subunits was confirmed in high glucose-treated podocytes and in glomeruli isolated from streptozotocin (STZ)-induced diabetic mice. Using live cell imaging, we illustrated that dynein-mediated post-endocytic sorting of nephrin was upregulated, resulting in accelerated nephrin degradation and disrupted nephrin recycling. In diabetic podocytopathy, *Dynll1* is one of the most upregulated dynein components that was recruited to endocytosed nephrin. This was corroborated by observing enhanced Dynll1-nephrin colocalization in podocytes of diabetic patients, as well as dynein-mediated trafficking and degradation of nephrin in STZ-induced diabetic mice. Knockdown of *Dynll1* attenuated lysosomal degradation of nephrin and promoted its recycling, suggesting the essential role of Dynll1 in dynein-mediated mistrafficking. Defining the role of dynein-mediated mistrafficking of nephrin in diabetes will not only fill the knowledge gap about the early events of DN, but also inspire novel therapeutics that target a broad spectrum of molecular events involved in the dynein-mediated trafficking.

**Translational Statement:** Diabetic nephropathy (DN), the leading cause of end stage kidney disease in the United States, is characterized by a podocytopathy with mistrafficking and depletion of the slit diaphragm protein nephrin, which in turn compromises the podocytes’ function in maintaining the glomerular filtration barrier. There is a critical need to define the trafficking mechanisms underlying the depletion of nephrin. Our work implicates cytoplasmic dynein, a trafficking complex that connects diabetes-triggered endocytosis with proteolytic pathways. Delineation of the dynein-driven pathogenesis of diabetic podocytopathy will inspire new therapies that potentially target a broad spectrum of molecules involved in dynein-mediated trafficking and degradation pathways.

## Introduction

Diabetic nephropathy (DN) is the most prevalent, acquired podocytopathy in humans and contributes to more than 50% of the cases of end-stage glomerulopathy^1^. Hyperglycemia and associated glomerular hyperperfusion are considered the major causes of DN; tight glucose control has clearly been shown to reduce the incidence of microalbuminuria and postpone the development of DN^2-4^. However, we lack a clear understanding of how hyperglycemia directly causes kidney injury, which in turn limits our ability to develop effective therapies.

Podocyte dysfunction is the hallmark of DN^5, 6^. Podocytes are highly differentiated, epithelial cells that encapsulate the glomerular capillary loops. The leading edge of podocytes branches into foot processes that interdigitate with those of neighboring podocytes to form slit diaphragms (SD), creating the molecular sieve of the glomerular filtration barrier (GFB). The SD consists of the extracellular domain of nephrin anchored in the cell membrane of the foot processes^7-9^. Microalbuminuria, an early presentation of DN, reflects an impaired GFB due to podocyte injuries that range from foot process effacement to more severe situation of podocyte detachment as illustrated by electron microscopy^10, 11^. In diabetic mouse models and in human DN, loss of nephrin and disruption of the normal SD architecture precedes the onset of microalbuminuria. This observation suggests that altered nephrin homeostasis influences the nanostructure of SDs during the initial phase of diabetic podocyte injury ^12, 13^. Studies using *in vivo* and *in vitro* models of DN observed enhanced nephrin endocytosis via multiple cell signaling pathways mediated by CIN85/RukL (Regulator of ubiquitous kinase/Cbl-interacting protein of 85 KD)^14^, protein kinase (PKC) α/β^15, 16^, PACSIN2 (PKC And Casein Kinase Substrate In Neurons 2) or p38 MAPK^17^. The key trafficking mechanism that is missing is the series of steps between the initial signaling-enhanced endocytosis described above and the outcome of nephrin depletion. Most of the studies on diabetes-induced mistrafficking have focused on the transient and short-distance endocytosis mediated by microfilaments^14-16, 18-21^, but the pathological processes following these initial endocytic events is unknown, leaving a critical knowledge gap, namely, how the diabetic milieu changes other, longer-range trafficking processes that are also involved in the homeostasis of the key SD proteins.

Kinesin and cytoplasmic dynein are motor proteins that guide reciprocal anterograde and retrograde transport of cargo proteins along the microtubule, between the surface membrane and the cytosol ^15^. The dynein transport complex consists of heavy chains, the ATPase subunit to generate energy for the sliding of the complex along microtubule^22^; light intermediate chains and intermediate chains that anchor cargo proteins or recruit adaptor proteins^23^; light chains that assemble and activate the whole complex^24, 25^; and dynactin 1, an required activator for the retrograde trafficking of dynein^23^. Dynein plays a unique role in modulating protein metabolism by mediating post-endocytic sorting of proteins and targeting them for ubiquitin-dependent degradation^22, 26-29^. The current study stemmed from two earlier findings. First, studies with streptozotocin (STZ)-induced diabetic rodents revealed increased dynein expression in retinal ganglion cells and skeletal muscle^30, 31^, supporting the idea that altered dynein-mediated trafficking might drive diabetic complications. Secondly, we recently described the pathogenesis of podocytopathy caused by mutations in the inverted formin 2 (*INF2*) gene, in which enhanced dynein-mediated trafficking depleted nephrin from the SD by diverting endocytosed nephrin into the lysosome degradation pathway^28^. Together, these findings illuminated a mechanism that may drive the development of podocytopathy in DN that we tested in the present study.

## Materials and Methods

The primers for real time quantitative PCR as well as the antibodies, siRNA duplex sequences, chemical compounds, and plasmids used in this study are listed in Tables 1 to 5, respectively.

**Table 1:**
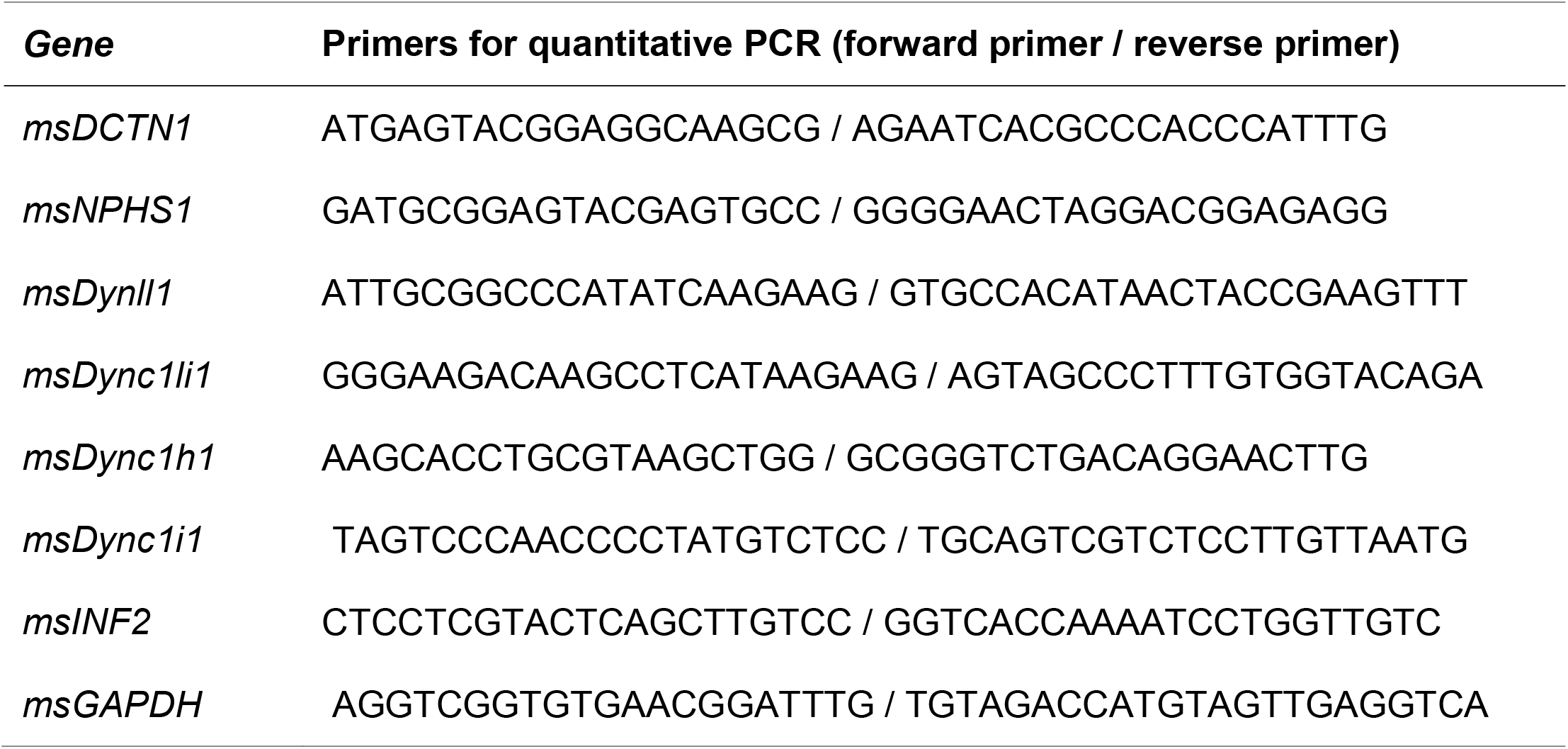
Primers for real time quantitative PCR

**Table 2:**
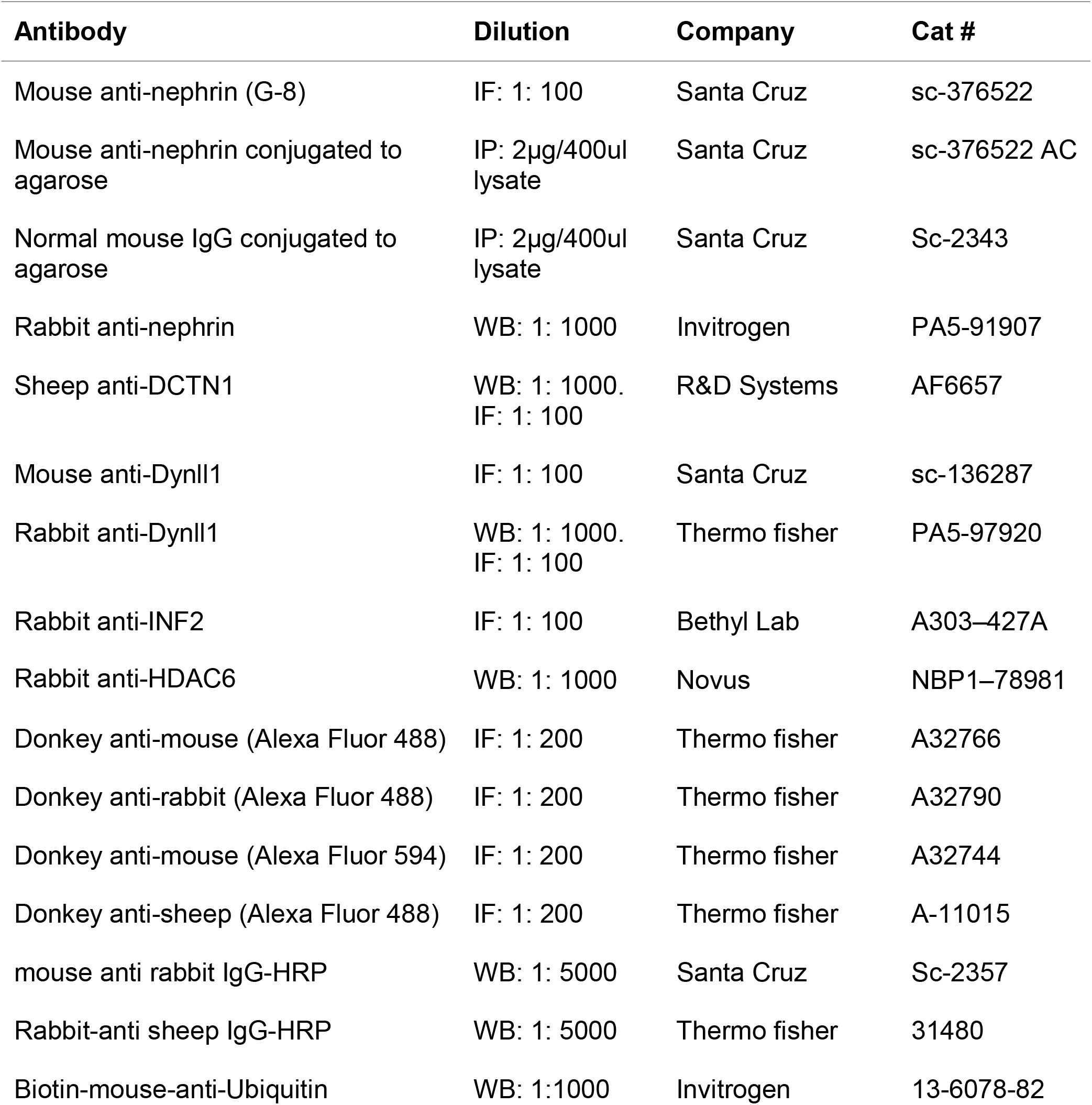
Antibodies [Immunofluorescent (IF), Western blotting (WB), Immunoprecipitation (IP)]

**Table 3.**
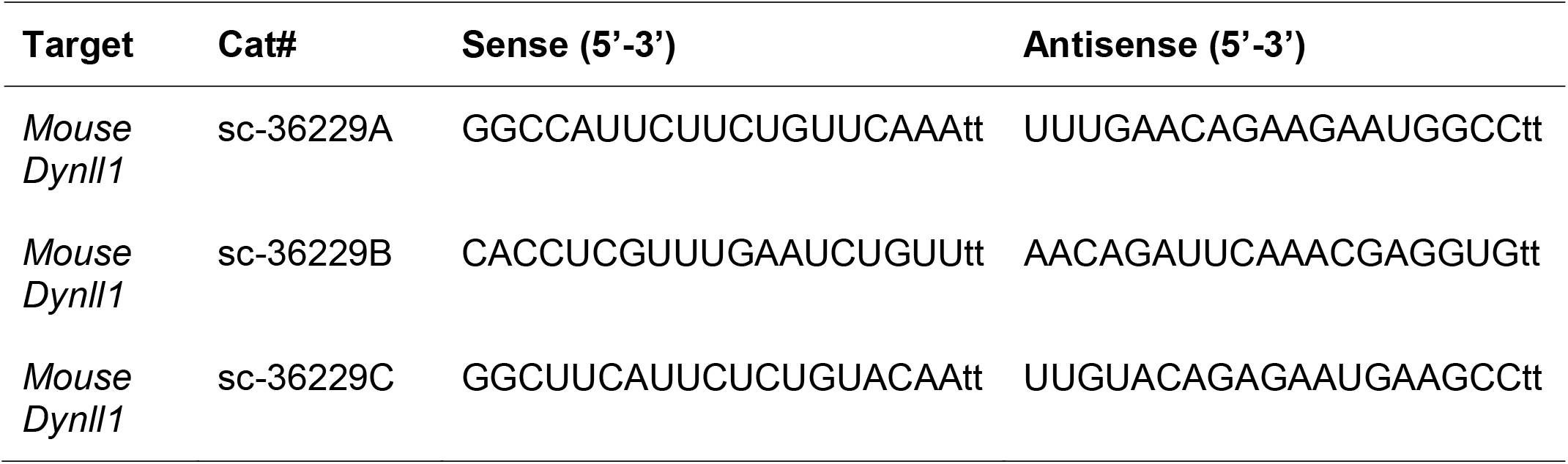
siRNA Duplex sequences

**Table 4:** Chemical compounds

| **Chemical compounds** | **Concentration** | **Company** | **Cat#** | **CAS#** |
| --- | --- | --- | --- | --- |
| Ciliobrevin D | 50 μM | Sigma | 250401 | 1370554-01-0 |
| Leupeptin | 200 μg/ml | MedChemExpress | HY-18234 | 103476-89-7 |
| Bortezomib | 100 nM | Sigma | 5.04314 | 179324-69-7 |
| Cycloheximide | 10 μM | Sigma | 01810 | 66-81-9 |
| Streptozotocin | 50 mg/kg, i.p.* daily x 5 | MedChemExpress | HY-13753 | 18883-66-4 |

\*Intraperitoneal injection

**Table 5:**
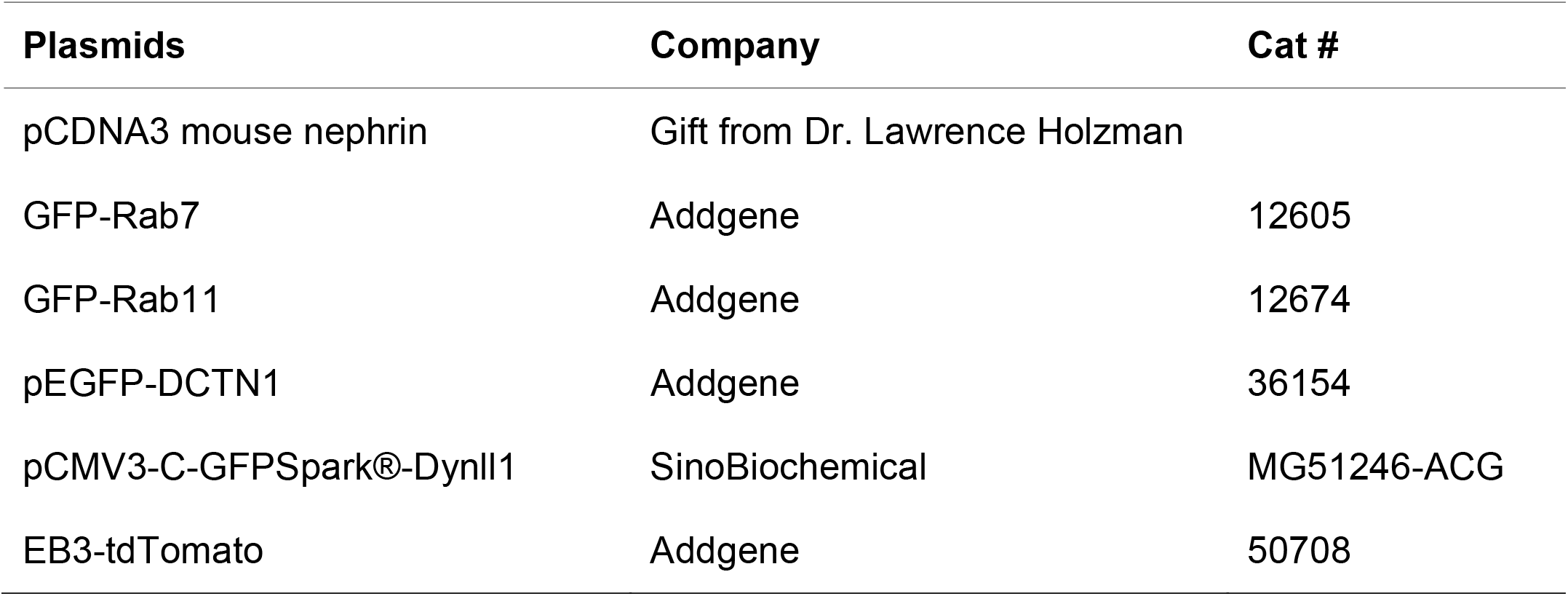
Plasmids

| Plasmids | Company | Cat # |
| --- | --- | --- |
| pCDNA3 mouse nephrin | Gift from Dr. Lawrence Holzman |  |
| GFP-Rab7 | Addgene | 12605 |
| GFP-Rab11 | Addgene | 12674 |
| pEGFP-DCTN1 | Addgene | 36154 |
| pCMV3-C-GFPSpark®-Dynll1 | SinoBiochemical | MG51246-ACG |
| EB3-tdTomato | Addgene | 50708 |

### Transcriptome analyses of dynein genes

Transcription analyses of cytoplasmic dynein genes were performed using *Nephroseq* kidney transcriptome databases and data-mining platform (www.nephroseq.org, University of Michigan, Ann Arbor, MI). Specifically, the transcription data for dynein components was retrieved from the Hodgin Diabetes Mouse Glom dataset (2013, Affymetrix Mouse Genome 430 2.0 Array platform)^32^. A cluster of overexpressed dynein genes were enriched in diabetic mice with a fasting glucose level >300 mg/dl (n=5), compared to mice with a fasting glucose level <300 mg/dl (n=16). In the analysis of the Woroniecka Human Diabetic Kidney Disease dataset (2011, Affymetrix Human Genome U133A 2.0 Array platform) ^33^ and the European renal cDNA bank (ERCB) nephrotic syndrome dataset (2018, RNA-Seq technique), a cluster of upregulated dynein genes were enriched in DN (n=10), compared to kidneys of healthy living donors (n=12).

In the Nephroseq data analysis platform, the two-sided *Student’s t-test* was used to compare the differential expression profiles of 2 groups. The *Log2 median centered expression* profile of the genes was expressed as a heatmap; the *Z score*s were normalized to depict relative values and color-coded (warmer colors represent higher relative expression levels and colder colors represent lower expression levels). The *Log2 (fold change, FC)* in expression of each individual gene in the kidney was compared between mice with fasting glucose >300 mg/dl and those with fasting glucose <300 mg/dl, and between human DN and healthy living donors. Furthermore, the *Pearson correlation* was used to analyze the relationship of individual dynein gene expression to the fasting glucose levels and albuminuria of diabetic mice in the Hodgin Diabetes Mouse Glom dataset (n=5 for the *db/db* mouse model, n=21 for all diabetic mouse models included in the study), or to the glomerular filtration rate (GFR) of human subjects in the Woroniecka Human Diabetic Kidney Disease dataset (n=9) and the ERCB Chronic Kidney Disease dataset (n=8).

### Cell culture and high glucose-induced podocytopathy

An immortalized mouse podocyte cell line^28^ which had been generated by expression of a temperature-sensitive mutant of the SV40 Large-T antigen per Saleem’s protocol^34^, was maintained in RPMI 1640 media with 10% FBS, 1% insulin–transferrin–selenium supplement (Gibco, Grand Island, NY), and 50 IU/ml penicillin/streptomycin. Podocytes were grown at 33°C and differentiated at 37°C for 2 weeks. The medium was replaced with RPMI1640 containing different concentrations of glucose, and mannitol was added to maintain the same osmolality among groups: 1) normal glucose (NG), glucose 5.5 mM+ mannitol 24.5 mM; 2) prediabetic level of glucose (PDB), glucose 11 mM+ mannitol 19 mM; and 3) high glucose (HG), glucose 30 mM alone. After 72 hours of treatment, cells were collected for total RNA extraction, cell biology assays, or biochemical experiments.

### Knockdown of Dynll1 in cultured podocytes

as previously described^28^, podocytes growing in a 6-well plate were transfected with 0.75µg (or 60 pmols) *msDynll1* siRNA Oligo Duplex (Santa Cruz #sc-36229, a pool of 3 mouse *Dynll1*-targeting siRNAs [A, B and C, 19-25 nt]), using the siRNA Transfection Reagent (Santa Cruz #sc-29528) according to the manufacturer’s instructions. Cells were used for experiments 72 hours after the transfection.

### Quantitative PCR

Total RNA was extracted from the podocytes or mouse glomeruli using the RNeasy Plus Mini Kit (Qiagen, #74134). 1 μg of RNA was reverse transcribed using Oligo dT primers and a high-capacity cDNA Synthesis Kit according to the manufacturer’s instructions (Applied Biosystems, #4368814). Quantitative PCR amplifications were performed using the Power SYBR Green PCR Master Mix (Applied Biosystems, #4368577) and an Applied Biosystem QuantStudio 7 Pro instrument. The primer sequences for real-time quantitative PCR were obtained from PrimeBank of Harvard University (https://pga.mgh.harvard.edu/primerbank/) and synthesized by Integrated DNA Technologies. The specificity of the primers was validated by the corresponding melting curves. Relative quantification of dynein gene transcription was performed using *GAPDH* as an endogenous control gene (constant expression across NG and HG conditions in cultured podocytes) ^35^and cells treated with NG as references. The *Log2 fold change* of each gene was calculated for comparison. For the *in vitro* study, the relative quantification of mRNA was compared among cells treated with different concentrations of glucose (5.5 mM, 11 mM, or 30 mM) and the assay was repeated in 3 independent experiments. For *in vivo* study, the relative quantification was compared between STZ-induced diabetic mice and vehicle-only controls (n=6 in each group).

### Co-Immunoprecipitation and Immunoblotting

Cultured podocytes were lysed on ice for 1 hour in Nonidet P40 lysis buffer with 0.5% deoxycholate, cOmplete™, Mini Protease Inhibitor Cocktail (Roche, #11836153001), and PhosSTOP phosphatase inhibitor cocktail (Sigma, #4906845001). For Co-Immunoprecipitation (Co-IP), cell lysates were immunoprecipitated with mouse-anti-nephrin conjugated agarose (Santa Cruz, sc-376522 AC, 2µg for 400μl cell lysate of 1×10^6^ cells), using normal mouse IgG conjugated agarose as control (Santa Cruz Biotechnology, sc-2343). For immunoblotting, the immunoprecipitated proteins or proteins in cell lysates were separated by SDS-PAGE and transferred to polyvinylidene fluoride or polyvinylidene difluoride (PVDF) membranes. After blocking, the blot was incubated with the primary antibody at 4°C overnight. After washes, the blot was incubated with the appropriate peroxidase-labeled-secondary antibody at room temperature for 30 minutes. Signals were detected using the Clean-Blot™ IP Detection Reagent (HRP) kit (Thermo Fisher, #21230) and SuperSignal West Dura Extended Duration Substrate (Thermo Fisher, #34075). For quantification, densitometry was performed using ImageJ software (NIH).

### Immunofluorescence Labeling of cultured podocytes

Podocytes grown on collagen I coated coverslips (Thermo fisher # A1142801) were fixed with 4% paraformaldehyde (Electron Microscopy Sciences, #30525-89-4) for 5 minutes, followed by standard labeling with primary and secondary antibodies as described in our previous study^28^.

### Immunofluorescent labeling of kidney sections

Paraffin-embedded kidney biopsy sections were obtained from the Pathology Department of the University of Iowa Hospitals and Clinics under an approved IRB protocol. Based on an established protocol^28^, the sections were deparaffinized and rehydrated. Antigen retrieval was performed by incubating the slides in IHC-Tek Epitope Retrieving buffer (IHC world, #IW-1000) in an IHC-Tek Epitope Retrieval Steamer (IHC world, #IW1102) for 45 minutes, followed by standard immunofluorescent staining. Dynll1 and nephrin were co-labelled by rabbit anti-Dynll1+ Alexa Fluor 488-conjugated Donkey anti-Rabbit IgG and mouse-anti nephrin+ Alexa Fluor 594-conjugated Donkey anti-mouse IgG. Fluorescence signals were captured from 3 non-sclerotic glomeruli of each section under a 63x oil immersion lens of an SP8 Leica confocal microscope. Dynll1-nephrin colocalization was mapped by using the *Colocalization threshold* plugin of Fiji software and quantified with the *Colo2* plugin as the *Pearson’s correlation coefficient* and the *Manders overlap coefficient*^36, 37^. These parameters were compared among DN (n= 30 glomeruli in 10 specimens), Focal and Segmental Glomerulosclerosis (FSGS, n= 26 glomeruli in 10 specimens), Minimal Change Disease (MCD, n= 17 glomeruli in 7 specimens) and normal kidney (n=9 glomeruli in 3 specimens).

### Nephrin Trafficking Model and Time-Lapse Imaging

As shown in the schematics in Figure 2A, cells transfected with the pcDNA3-nephrin plasmid [Gift from Dr. Holzman ^7^] were incubated with a mouse anti-nephrin antibody recognizing the extracellular domain (Santa Cruz #sc-376522), followed by an Alexa Fluor 594-conjugated donkey anti-mouse IgG (Thermo fisher #A32744) to trigger crosslinking and endocytosis of nephrin. As described in our previous study^28^, live cell imaging was performed by scanning cells every 5 seconds under a Leica SP8 Confocal microscope for a total of 20 minutes. Single molecule tracking of endocytosed nephrin with the co-expressed GFP-Dynll1 fusion protein (Sino Biological, pCMV3-C-GFPSpark-Dynll1, # MG51246-ACG), EGFP-DCTN1 fusion protein (Addgene pEGFP-p150Glued, #36154) or tdTomato-EB3 fusion protein (Addgene, EB3-tdTomato, #50708) was performed as 5 repeated measurements × 5 cells × 3 independent experiments, and the data analysis was performed using the Fiji plugin KymographClear coupled with KymographDirect. The fraction and the average velocity of the retrograde (red), anterograde (green), and static (blue) trafficking events were calculated and compared in cells with different treatments.

### Surface biotinylation–based recycling assay

As described in our previous work^28^ and as outlined in schematics in Figure 4C, (1) podocytes transfected with the pcDNA3-nephrin plasmid were incubated on ice in EZ link sulfo-NHS-SS-biotin (Pierce, #21331) for 45 minutes. (2) Cells were then returned to 37°C and underwent endocytosis, followed by (3) surface stripping by incubating with 100 mM Mesna (Santa Cruz, CAS 19767-45-4) at 4°C to remove biotin from uninternalized nephrin. (4) Cells were returned to 37°C to allow for recycling of the endocytosed nephrin for 2 hours, followed by (5) a second surface stripping with Mesna to remove biotin from the recycled nephrin (R, harvested after step 4). Cells without the second Mesna strip served as a protein degradation control (DC, harvested after step 4). The biotinylated nephrin in cell lysates was pulled down by incubation with streptavidin-conjugated beads (Pierce, #20353) at 4°C overnight with continuous rotation, and the biotinylated nephrin was measured by immunoblotting using anti-nephrin. The percentages of recycled nephrin were calculated as [(degradation control, DC) − (recycling, R)]/ (degradation control, DC)]x100%.

### Streptozotocin-induced diabetic nephropathy in C57BL/6J mice

As described in a published protocol ^*38, 39*^, 8-week-old, fasted C57BL/6J wild-type mice (male: female ratio = 1:1) received i.p. injection of streptozotocin (STZ, MedChemExpress LLC, # HY-13753) dissolved in 0.1□M sodium citrate buffer (pH = 4.5), at the dose of 50□mg/kg body weight, daily for 5 consecutive days. Age- and gender-matched vehicle control mice received i.p. (intraperitoneal) injections of sodium citrate buffer of the same volume and regimen. Whole blood glucose was measured in tail vein blood specimens using the Germaine™ Laboratories AimStrip ™ Plus Blood Glucose Testing System. Urine albumin and creatinine were measured in specimens collected before and at different months after the injections, using the Albumin ELISA Quantitation kit (Bethyl Laboratories Inc.) and a colorimetric creatinine quantification kit (Bioassay Systems). The urine albumin-to-creatinine ratio (ACR) was calculated for comparison.

### Kidney specimen handling, histology, and quantification

Mice were anesthetized by isoflurane inhalation and then perfused with EZ link sulfo-NHS-SS-biotin per an *in vivo* surface biotinylation protocol before sacrifice^40^. The mouse kidneys were then harvested for light microscopy (LM), transmission electron microscopy (TEM), immunogold labeling (IGL), and for molecular biology studies at the Comparative Pathology Laboratory and the Central Microscopy Research Facility (CMRF) at the University of Iowa. The glomerular hypertrophy was analyzed by quantifying the area of each individual glomerulus. Podocytopathy was quantified by measuring the percentages of capillary loops covered by injured podocytes with effaced foot processes or disassembled SDs. For immunogold labeling^28^, 8 nm sections were placed on carbon-coated and glow-discharged, formvar-coated nickel slot grids; blocked grids were incubated in a mouse anti-podocin antibody (Sigma Aldrich, #SAB4200810) or a rabbit anti-nephrin antibody (Invitrogen, # PA5-91907) at room temperature, followed by a goat anti-rabbit Au 12 nm conjugated (Jackson ImmunoResearch Laboratories Inc. #111-205-144) and a goat anti-mouse Au 6 nm conjugated (Jackson ImmunoResearch Laboratories Inc. #115-195-146). Mouse glomeruli were isolated based on the standard protocol as described in reference^41^, and then lysed in RLT buffer (RNeasy Plus Universal Mini Kit, Qiagen, # 74034) or Nonidet P40 lysis buffer for mRNA and protein analysis, respectively.

### Statistical Analyses

Data analyses were performed using SPSS version 10.0. Immunofluorescent signal analysis and densitometric analysis of immunoblots were done using National Institutes of Health (NIH) Fiji/ImageJ software. Data were expressed as mean ± SEM. An independent sample *t* test was used to compare the difference between two groups. *One-way ANOVA* was used for comparisons among multiple groups, and a post-hoc *q* test was used to compare the difference between groups. In a two-tailed test, *p*<0.05 was considered significant.

### Study Approval

Paraffin-embedded kidney biopsy sections were obtained from the Pathology Department of the University of Iowa, and the University of Iowa Institutional Review Boards approved the secondary use of archived kidney biopsy specimens. The animal protocol was approved by the Institutional Animal Care and Use Committee of the University of Iowa and is in accordance with the National Institutes of Health guidelines for use of live animals. All work was carried out in accordance with the principles and procedures outlined in the NIH guidelines.

## Results

### 1. Dynein expression is upregulated in DN and podocytes with hyperglycemic injury

We re-analyzed existing transcriptome datasets of DN in human and mouse models of diabetes. The Nephroseq datasets revealed increased expression of dynein components that positively correlated with fasting glucose levels in diabetic mouse models (**Figure 1A**) and with the decline of glomerular filtration rate (GFR) in human DN (**Figure 1B**).

**Figure 1.**
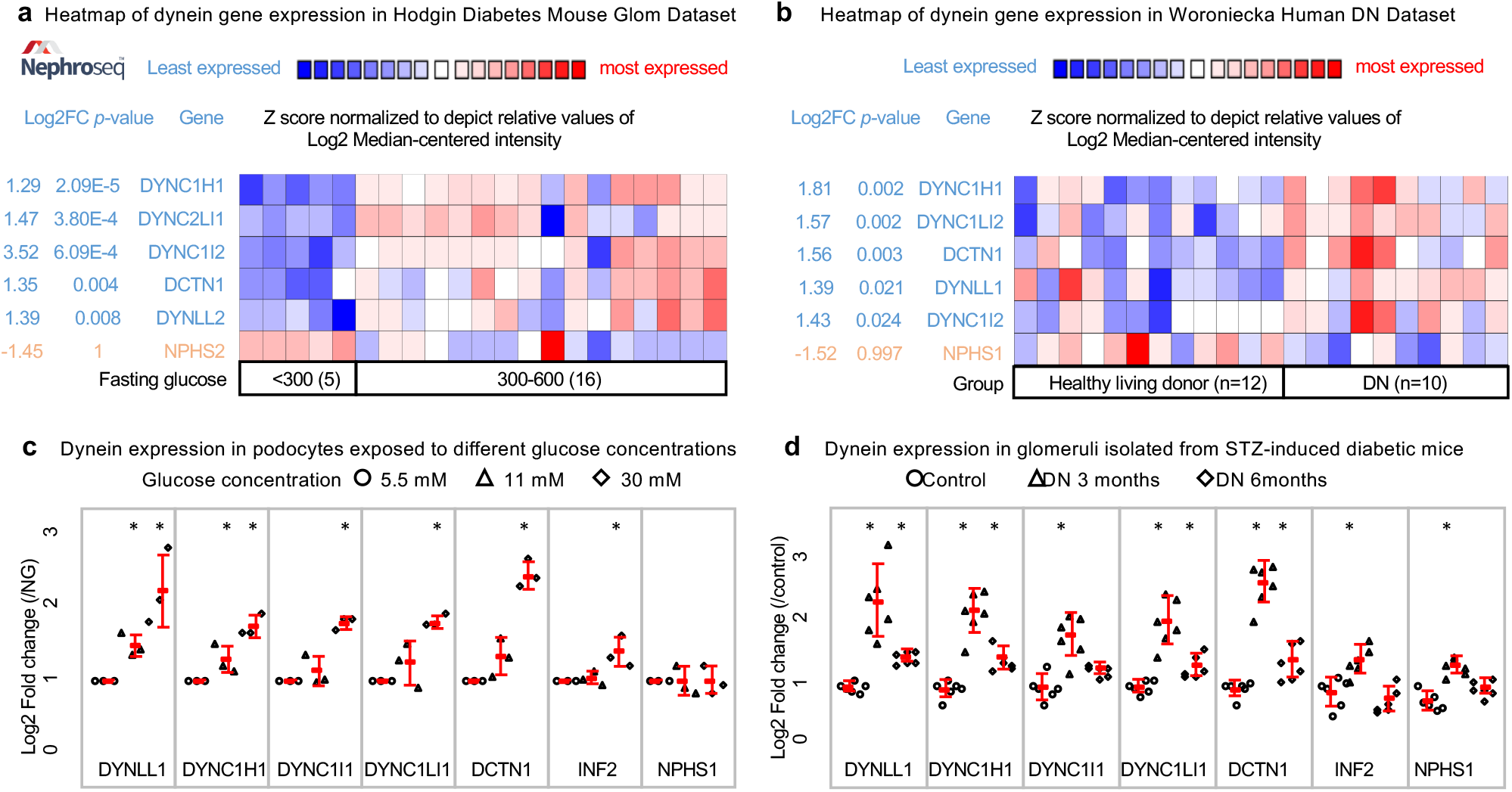
Figure 1. Increased dynein expression in diabetic nephropathy. a: Transcription analyses of Hodgin Diabetes Mouse Glom dataset in Nephroseq revealed upregulated dynein expression correlated with hyperglycemia in diabetic mice. Heatmap shows a cluster of dynein genes with increased expression in glomeruli of diabetic mice with fasting glucose >300 mg/dl. The Log2 median centered expression level of each individual dynein gene is color coded, reflecting *Z* scores normalized to depict relative value. b: Transcription analyses in Woroniecka Human Diabetic Kidney Disease dataset revealed upregulated dynein expression correlated with decline of GFR in human diabetic nephropathy. Heatmap shows a cluster of dynein genes with increased expression in human diabetic nephropathy. c & d: Quantification of mRNA levels of dynein components in podocytes growing in media containing different glucose concentrations [C. HG: high glucose, 30 mM; PDB: pre-diabetic level of glucose, 11 mM; and NG: normal glucose, 5.5 mM; n=3 independent experiments, \**p<0*.*05 vs. NG*] or in glomeruli isolated from STZ-induced diabetic mice (D) were measured by relative quantitative PCR (normalized to *GAPDH*) (D, n=6 mice each group. \**p<0*.*05 vs. vehicle control*)

In the Hodgin Diabetes Mouse Glom dataset^32^, we found a cluster of dynein genes with increased expression in diabetic mice with fasting glucose >300 mg/dl, compared to those with fasting glucose <300 mg/dl (**Figure 1A**). This cluster included *DYNC1H1* (dynein cytoplasmic 1 heavy chain 1), *DYNC2LI1* (dynein cytoplasmic 2 light intermediate chain 1), *DYNC1I2* (dynein cytoplasmic 1 intermediate chain 2), *DCTN1* (dynactin subunit 1) and *DYNLL2* (dynein light chain LC8-type 2). In *db/db* mice, the kidney expression of *DYNC1LI1* (*r*^*2*^=0.856) and *DYNC1LI2* (*r*^*2*^=0.870) correlated with their fasting glucose levels. In all diabetic mice that were analyzed, the kidney expressions of *DCTN1* (*r*^*2*^=0.355), *DYNC1H1* (*r*^*2*^=0.676), *DYNC2LI1* (*r*^*2*^=0.401) and *DYNC1I1* (*r*^*2*^=0.549) correlated positively with fasting glucose levels (**Supplementary Figure 1 B**). Additionally, the expression of *DYNC1LI2 (r*^*2*^*=0*.*612), DYNC1I1 (r*^*2*^*=0*.*898)* and DYNLT3 (Dynein Light Chain Tctex-Type 3) *(r*^*2*^*=0*.*599)* correlated with the severity of albuminuria in *eNOS-/-db/db* mice and STZ-induced DN models (**Supplementary Figure 1A**).

Coordinate expression of dynein genes was also found in the Woroniecka Human Diabetic Kidney Disease dataset^33^ (**Figure 1B**). *T*he expression level of *DYNC1I1 (r*^*2*^=0.535*) and DYNC2LI1* (*r*^*2*^=0.654) correlated inversely with GFR in human DN. In the ERCB Chronic Kidney Disease dataset, expression levels of *Dynll1* (*r*^*2*^=0.919), *DCTN1* (*r*^*2*^=0.649) correlated inversely with the GFR in human DN (**Supplementary Figure 1C**).

We next compared the mRNA levels of dynein components in cultured podocytes growing in media containing different glucose concentrations and in glomeruli isolated from the STZ-induced diabetic mice. Compared to cells growing in NG (glucose 5.5 mM with 24.5 mM of mannitol to maintain the same osmolality), cells growing in PDB (glucose 11 mM with 19 mM of mannitol), or HG (glucose 30 Mm alone) showed increased expression of dynein components (**Figure 1C**). Among them, *Dynll1 and DCTN1* had the most elevated expression in response to high glucose exposure, suggesting they are potential direct responders to hyperglycemia. Expression of the dynein components was increased and peaked in glomeruli isolated from diabetic mice three months after STZ injection (**Figure 1D**), when the mice have developed significant hyperglycemia >300 mg/dl (Figure 5D). The expression of *INF2*, which encodes a sequestrator for Dynll1, was only slightly increased in HG-treated cells and in the early stage of STZ-induced DN (three months) but was far less prominent than that of *Dynll1*. The expression of *NPHS1*, the gene encoding nephrin, was not reduced in diabetic podocytopathy *in vitro* or *in vivo*; instead, its expression increased in the early stage of STZ-induced DN.

### 2. Dynein-mediated post-endocytic sorting of nephrin is enhanced in high glucose-induced podocyte injury

We found that high glucose treatment of podocytes increased the dynein-dependent trafficking of nephrin to the lysosome. Here, we used an *in vitro* model of nephrin endocytosis triggered by antibody-mediated crosslinking ^28^ (**Figure 2A**). The recruitment of dynein components to endocytosed nephrin was measured by a co-immunoprecipitation assay (**Figure 2B**) and visualized with an immunofluorescence colocalization assay (**Figure 2C**). Growing podocytes under high glucose conditions (HG, 30mM) resulted in higher levels of Dynll1 and DCTN1 recruitment to the endocytosed nephrin than observed in podocytes grown under normal glucose concentrations (NG, 5.5). Additionally, we found increased recruitment of Histone deacetylase 6 (HDAC6) to endocytosed nephrin in HG-treated cells. HDAC6 is the protein that bridges the dynein transport complex with ubiquitinated protein that is designated for degradation ^42, 43^. The recruitment of both dynein components and HDAC6 was diminished by antagonizing dynein with the addition of Ciliobrevin D (50µM, a direct dynein ATPase inhibitor)^44^ to the media or by siRNA-mediated knockdown of *Dynll1*, a component essential for the integrity of the dynein transport complex^45^. These data suggest HDAC-6 is recruited via dynein, consistent with previous studies showing the association of Dynein with HDAC6.^46^

**Figure 2.**
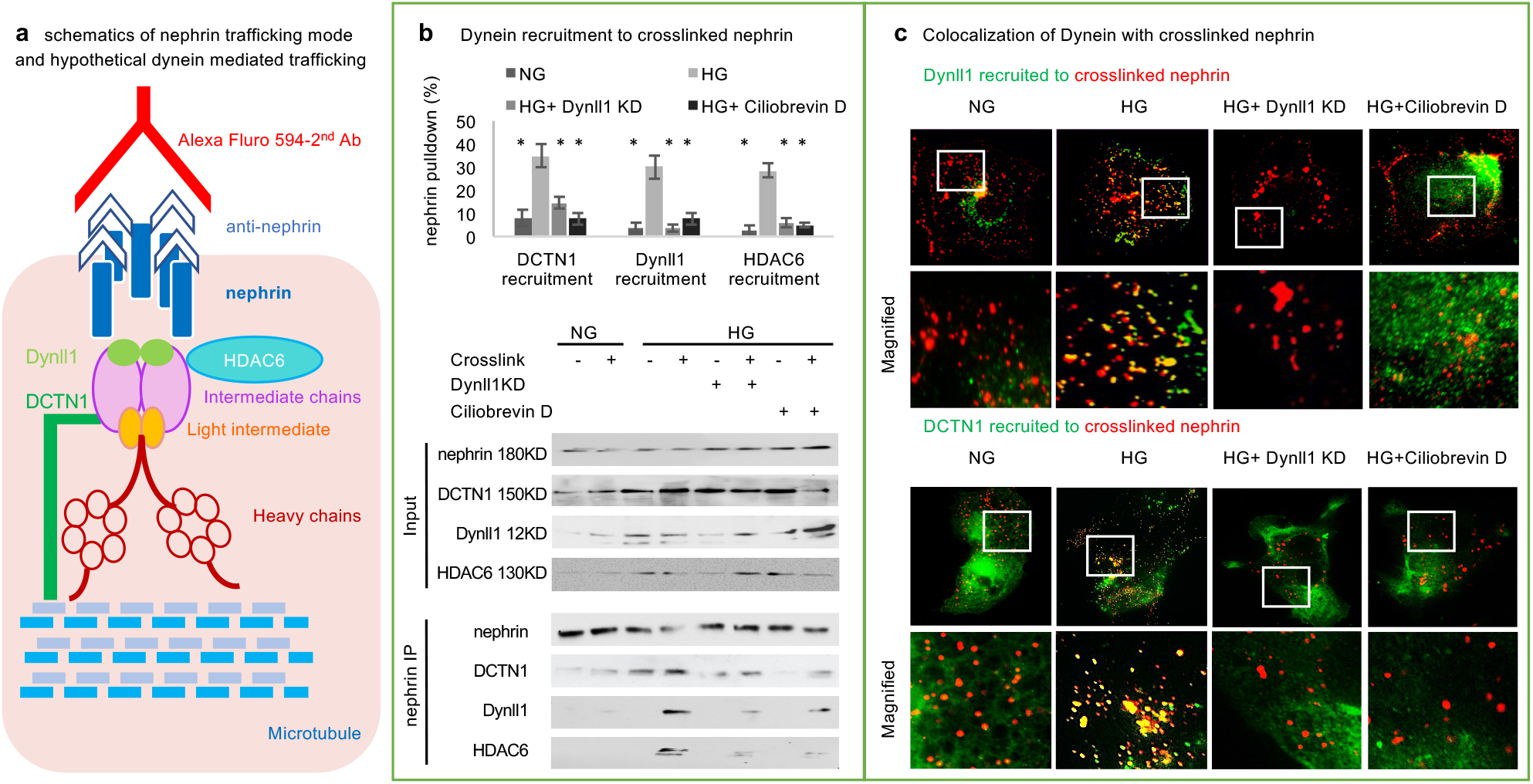
Enhanced dynein-mediated post-endocytic trafficking of nephrin in hyperglycemic conditions. a. *In vitro* nephrin trafficking assay in mouse podocytes using antibody-mediated crosslinking of nephrin: nephrin molecules expressed on the podocyte surface were crosslinked by an anti-nephrin primary antibody and Alexa Fluro 594 labeled secondary antibody, which triggered nephrin endocytosis and recruitment of trafficking adaptors. As demonstrated by Co-IP (b, by measuring dynein components in nephrin pulldown) and an immunofluorescence colocalization assay (c, crosslinked-nephrin labeled in red by Alexa Fluor 594, Dynll1 and DCTN1 labeled in green by Alexa Fluor 488), high glucose treatment (HG, 30 mM) increased the recruitment of Dynll1, DCTN1 and HDAC6 to crosslinked nephrin, compared to normal glucose (NG, 5.5 mM). These changes could be diminished by inhibiting dynein activity (using Ciliobrevin D, 50 µM) or by knocking down *Dynll1 (Dynll1 KD)* in podocytes. (n=3, *\*p<0*.*05 vs*. HG).

### 3. Dynein enhanced the retrograde trafficking of nephrin in high glucose treated podocytes

We imaged live cells to track endocytosed nephrin with co-expressed GFP-Dynll1, EGFP-DCTN1 or tdTomato-End-binding protein (EB3). As shown in **Figure 3A**, kymograph analysis demonstrates that the endocytosed nephrin shared the same trajectory with tagged Dynll1 and DCTN1, and the nephrin moved in the opposite direction along the microtubule to tdTomato-EB3, a marker for the plus-ends of microtubules ^47^. These findings indicate a dynein-mediated retrograde trafficking of endocytosed nephrin away from the plus ends.

**Figure 3.**
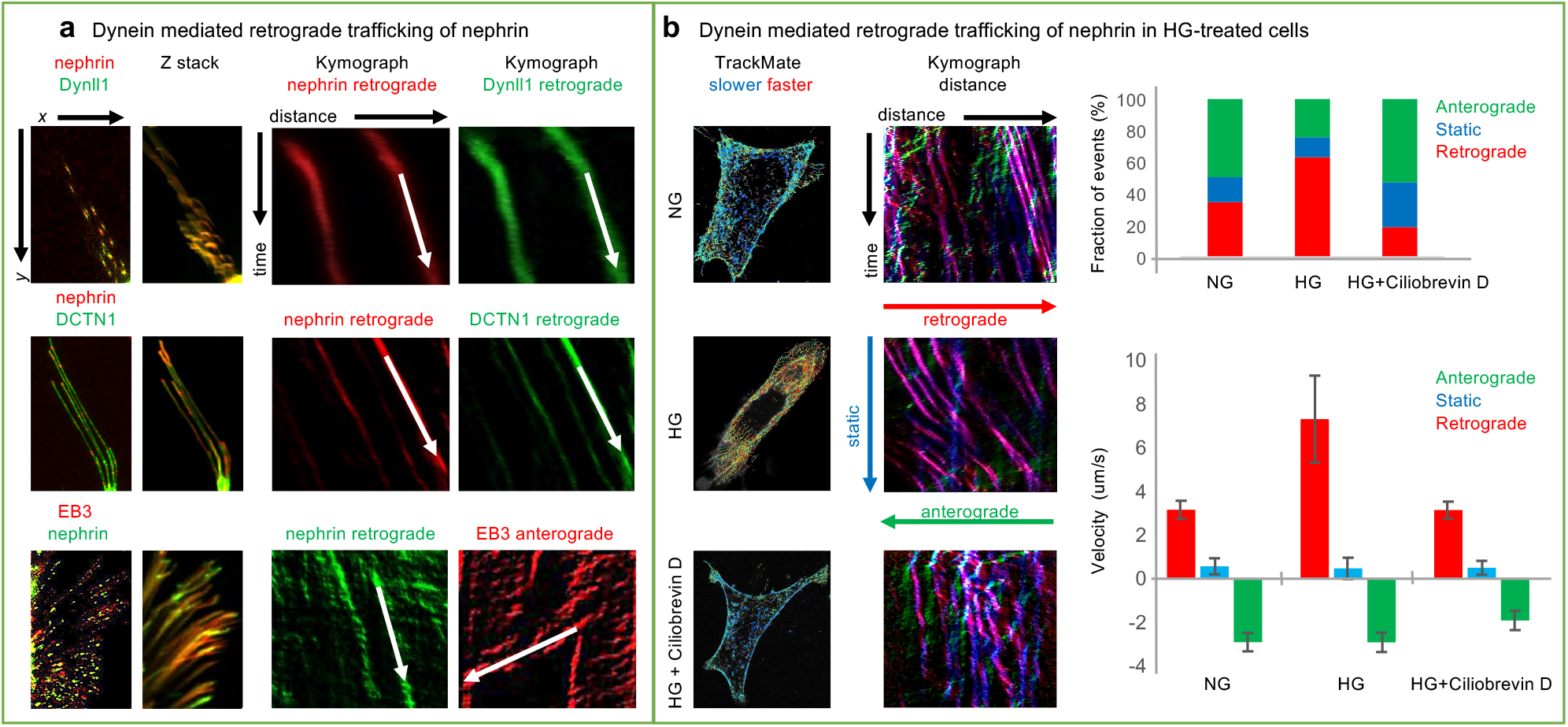
Live cell imaging of dynein-mediated trafficking. a. Trajectory analysis of nephrin with co-expressed GFP-Dynll1, GFP-DCTN1 or tdTomato-EB3 by using live cell imaging and kymograph analysis. Nephrin and dynein components (Dynll1 or DCTN1) shared the same trajectory of retrograde trafficking to the minus-end of microtubules. The track of nephrin was parallel to but moving in the opposite direction to that of EB3, which labels the plus-ends of microtubules. b. The post-endocytic trafficking of nephrin was compared in podocytes growing in NG, HG, or HG in the presence of Ciliobrevin D (50 µm). The tracks of nephrin particles were illustrated by using *TrackMate* of Fiji software, the warm color reflecting a higher velocity and the cooler color a lower velocity. The fractions and velocities of anterograde (green, from cytosol to surface membrane), retrograde (red, from surface membrane to cytosol) and static (blue) trafficking events of nephrin in 5 repeated measurements × 5 cells × 3 independent experiments were collected and quantified by using *KymographClear* coupled with *KymographDirect* software.

We then compared the trafficking events in podocytes growing in NG (5.5 mM) or HG (30 mM). As shown in **Figure 3B**, in HG treated cells, there were significantly more and faster retrograde events (from surface membrane to cytosol), as reflected by increased fractions and velocities of retrograde events (highlighted in red) and reduced anterograde events, i.e., recycling from cytosol to surface membrane (highlighted in green), compared to cells growing in NG. These differences were suppressed by incubating cells with Ciliobrevin D, indicating an enhanced dynein-dependent post-endocytic trafficking in our *in vitro* model of diabetic podocytopathy.

### 4. Dynein-mediated trafficking alters nephrin degradation and recycling in high glucose treated podocytes

We observed increased sorting of nephrin to the late endosome/lysosomes, which is marked by Rab7 in HG-treated podocytes (F**igure 4A**). This is consistent with increased dynein recruitment that would be expected to direct endosomes towards lysosomes for degradation. To test this idea, we followed the degradation of endocytosed nephrin in HG-treated podocytes using a cycloheximide chase to inhibit synthesis of new nephrin. Over this period, there was a dramatic loss of nephrin in HG-treated podocytes that was significantly greater than the loss in NG-grown podocytes (**Figure 4B**). Moreover, degradation of nephrin was suppressed by Leupeptin, an inhibitor of lysosomal proteases. Finally, it was also blocked by Ciliobrevin D, suggesting that HG-induced degradation of nephrin was dynein-dependent.

**Figure 4.**
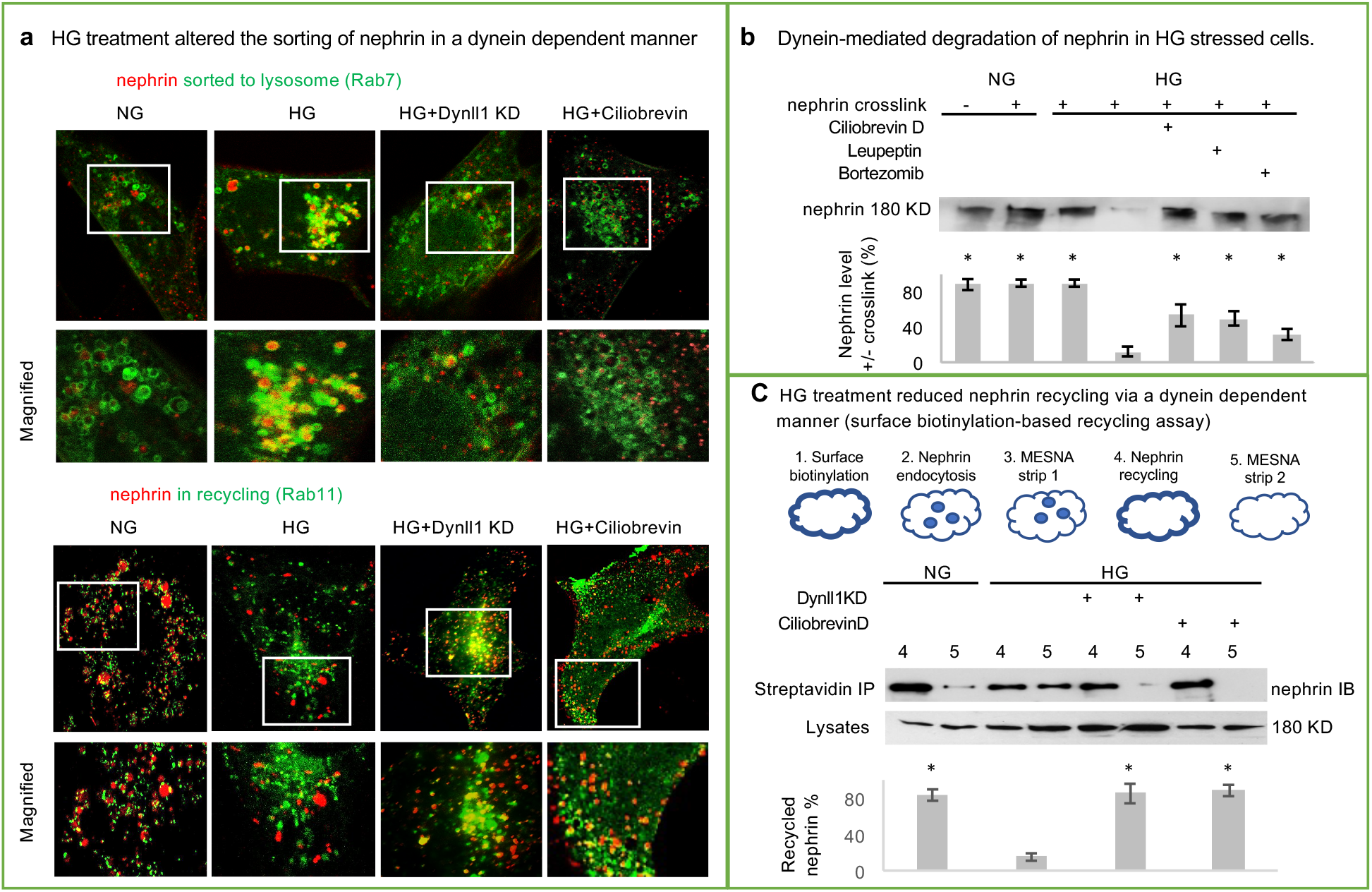
Dynein-mediated nephrin degradation and recycling in HG-treatment podocytesa. a. Tracking of nephrin and the co-expressed GFP-Rab7 or GFP-Rab11. Endocytosed nephrin proteins in HG-treated cells were enriched in Rab7-positive lysosomes with reduced Rab11 coating, as compared to NG-treated cells. These changes were much less pronounced in HG-treated cells in which dynein activity was inhibited with Ciliobrevin D or *Dynll1* was knocked down (*Dynll1 KD*) with siRNA. b. Cells were pretreated with cycloheximide (to reduce the background of newly synthesized nephrin) and underwent nephrin crosslinking and endocytosis. The extent of post-endocytic degradation of nephrin in these cells was reflected by the reduction in nephrin protein levels as measured by Western blotting. Nephrin crosslinking in HG-treated cells resulted in reduced nephrin (normalized to cells without crosslink x 100%), which was rescued by incubation with Ciliobrevin D (50µM), leupeptin (200 µg/ml) or Bortezomib (100 nm). n=3, *p<0*.*05 vs*. HG with nephrin crosslinking. c. Nephrin recycling was examined by using a surface biotinylation-based recycling assay as outlined in the schematics. Cells went through the following steps 1) biotinylation of all surface nephrin, 2) antibody-mediated internalization of nephrin, 3) first surface stripping with Mesna to remove biotin, i.e., to leave only the internalized nephrin biotinylated (total endocytosed), 4) recycling of the biotinylated nephrin (degradation control, DC), 5) second surface stripping with Mesna to remove biotin from the recycled nephrin (recycling, R). Cell lysates collected at steps 4 and 5 were subjected to streptavidin IP to pull down biotinylated nephrin, and the percentages of recycled nephrin were determined with the formula: % recycled = [(degradation control, DC) − (recycling, R)]/ (degradation control, DC)]x100%. *\* p<0*.*05 vs*. HG, n=3.

Next, we tested whether the enhanced dynein-mediated trafficking and degradation seen in HG-treated cells is associated with decreased recycling of nephrin. As shown in **Figure 4A**, by tracking endocytosed nephrin with coexpressed GFP-Rab11, a marker for recycling endosomes, we found a reduced degree of nephrin recycling in cells with exposure to HG. Further, using a surface biotinylation-based recycling assay (described in **Figure 4C**), we showed that internalized nephrin primarily recycled back to the cell surface in podocytes growing in NG, but that the recycling was greatly diminished under hyperglycemic conditions. Nephrin recycling in the HG-treated cells was restored with the addition of Ciliobrevin D or knockdown of *Dynll1*, supporting the essential role for dynein, especially Dynll1, in mediating the altered trafficking of nephrin in podocytes under hyperglycemic stress.

### 5. Dynein-mediated mistrafficking of nephrin related to its homeostasis *in vivo*

We used the STZ-induced diabetic mice as an *in vivo* model to determine whether dynein-mediated triage and homeostasis of nephrin observed in HG-treated podocytes *in vitro* correlate with the onset of nephropathy. STZ-induced diabetic mice developed significant hyperglycemia > 300mg/dl after two months (**Figure 5D**), and albuminuria after three months (**Figure 5E**). By month three, these diabetic mice developed histologic features of DN, including mesangial expansion and glomerular hypertrophy (reflected by increased size of glomeruli, **Figure 5F**).

**Figure 5.**
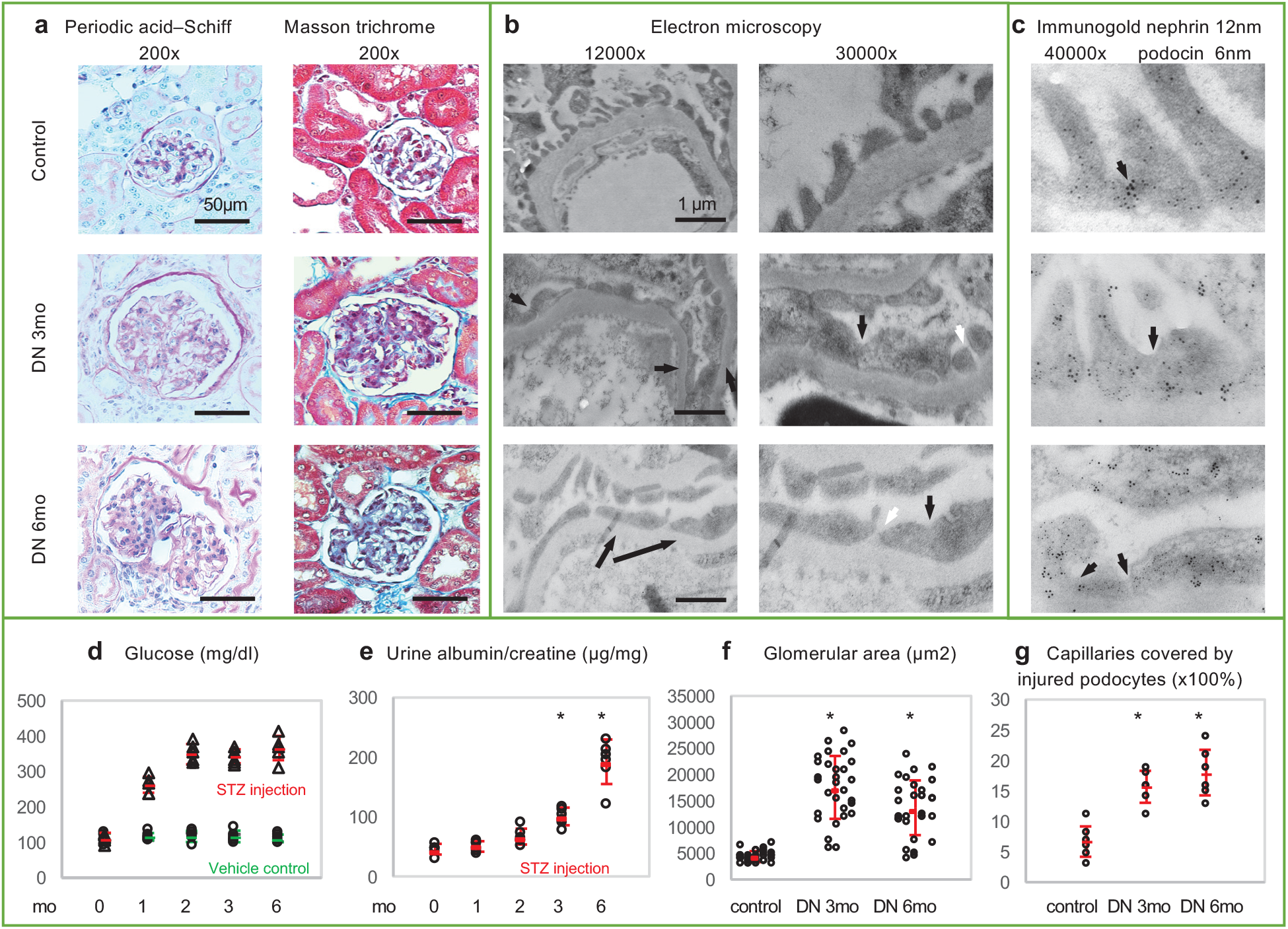
Podocytopathy in streptozotocin induced diabetic mice. The histological and ultrastructural features of diabetic glomerulopathy and podocytopathy developed in mice three months after receiving i.p. injection of *s*treptozotocin, as shown by periodic acid-Schiff staining and Masson Trichrome staining (a, Scale Bars = 50µm) and transmission electron microscopy (b,Scale Bars = 1µm). In mice with STZ-mediated DN, podocytopathy presented with effacement (black arrows) of foot processes, or cortical dislocation of slit diaphragms (white arrows). Podocytes with STZ-induced diabetic injury are characterized by the dislocation of nephrin and podocin from the slit diaphragm to the cytosol (arrows) as detected by immunogold labeling (c). Hyperglycemia (d) and albuminuria (e), quantified as the albumin/creatinine ratio, in mice with STZ-mediated DN (*\*p<0*.*05 vs*. pre injection, n=6). Histology and ultrastructural features of diabetic glomerulopathy were expressed as the average glomerular area (f, n=6 mice x 5 glomeruli per mouse) and the percentage of glomerular capillaries covered by effaced foot processes (g, n=6). *\*p<0*.*05 vs*. vehicle control.

The transmission electron microscopy (TEM) demonstrated features of podocytopathy reflected by effacement of foot process (**Figure 5B**), cortical relocation of SDs (white arrows), and an increased percentage of capillaries covered by abnormal podocytes (**Figure 5G**). By immunogold labeling, we showed the “trapping” of nephrin and podocin in the cytosol of podocytes in the STZ-induced diabetic mice, instead of localizing to the sites of SDs, as seen in the control mice (**Figure 5C**).

Immunofluorescence labelling of kidney sections from STZ-induced diabetic mice revealed overexpressed Dynll1 along the glomerular capillaries that colocalized with nephrin (**Figure 6A**). As depicted by *Manders overlap coefficients*, the colocalization of Dynll1 with nephrin was significantly higher in podocytes of diabetic mice. This was corroborated biochemically with coimmunoprecipitation experiments showing increased recruitment of DCTN1 and Dynll1 to nephrin from glomeruli isolated from STZ-induced diabetic mice (**Figure 6B**), as we had seen in cultured podocytes (**Figure 2B**). This corresponded to the reduced amount of surface nephrin--as measured in streptavidin pulldown—following the *in vivo* surface biotinylation assay^48^. We also observed reduced nephrin protein but not mRNA levels (**Figure 1D**) as well as nephrin ubiquitination in diabetic kidneys (**Figure 6B**), suggesting a dynein-mediated trafficking of nephrin that promotes its ubiquitin-mediated degradation in lysosomes.

**Figure 6.**
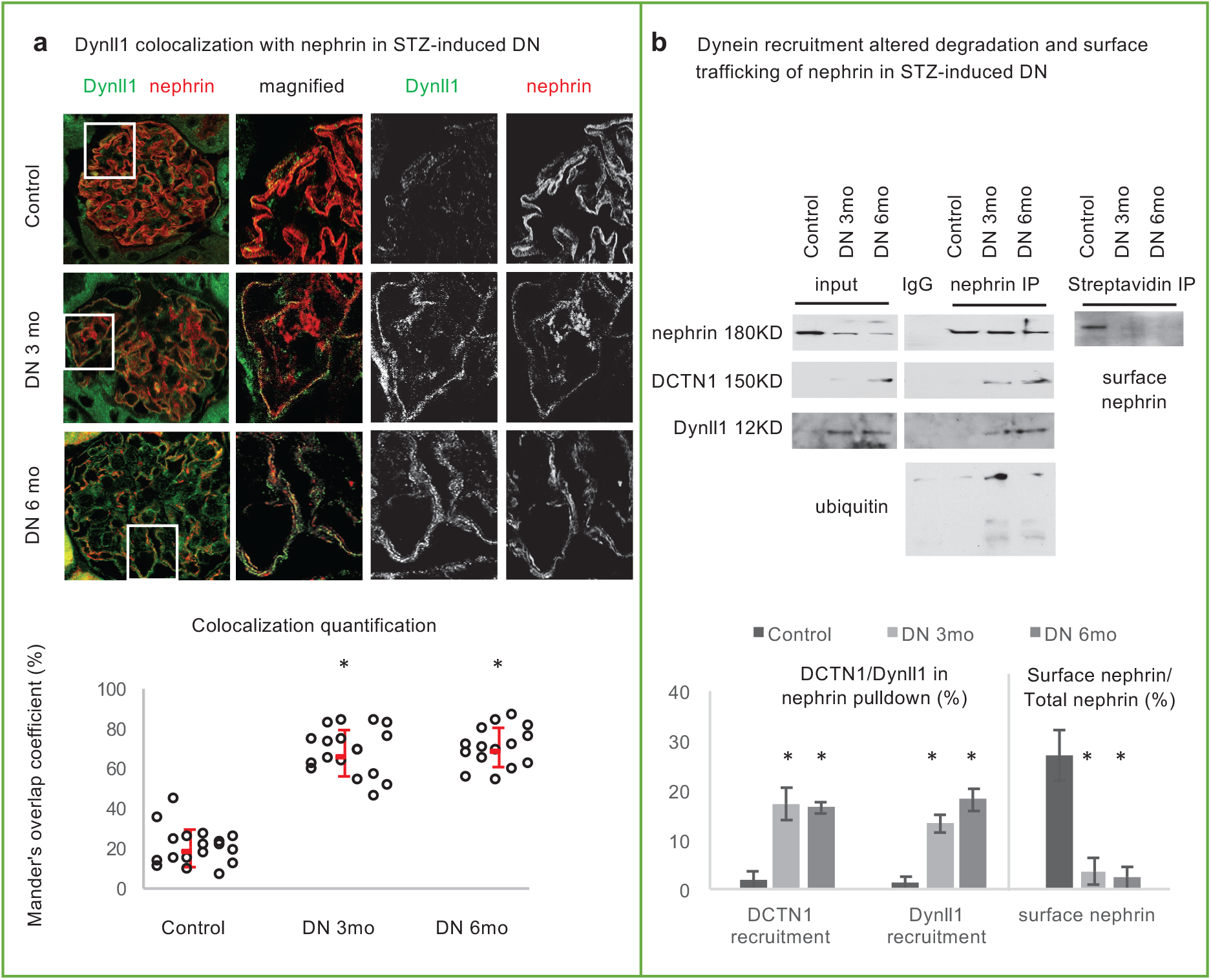
In vivo evidence of dynein related mistrafficking and degradation of nephrin. a. Immunofluorescent staining showed that overexpressed Dynll1 co-localizes with nephrin in STZ-induced mouse diabetic podocytopathy. The amount of Dynll1 colocalized with nephrin in non-sclerotic glomeruli was quantified using the *Colo2* plugin of Fiji software using the *Manders overlap coefficient* for comparison. *\*p<0*.*05 vs*. vehicle control, n= 6 mice x 3 glomeruli per mouse. b. The recruitment of Dynll1 and DCTN1 was examined in nephrin pulldown and were expressed as the ratio to total nephrin, reflecting the involvement of dynein in nephrin trafficking *in vivo*. Nephrin protein targeted for ubiquitin-mediated degradation was examined by measuring ubiquitinated nephrin (IP with anti-nephrin and IB with anti-ubiquitin). Nephrin localized on the podocyte surface was measured by streptavidin IP following the *in vivo* surface biotinylation assay, expressed as the percentage of total nephrin for comparison. *\*p<0*.*05 vs*. vehicle control, n= 6.

### 6. Evidence of dynein-mediated trafficking of nephrin in human DN

Upregulated Dynll1 expression co-localizing with nephrin was recently recognized as a feature of enhanced dynein-mediated trafficking of nephrin *in vivo* in a form of FSGS caused by pathogenic mutations in *INF2*^28^. In podocytes harboring INF2-R218Q mutation, the R218 mutant of INF2 was found to dissociate Dynll1, release it for nephrin recruitment and subsequently facilitate the dynein-mediated post-endocytic sorting of nephrin. Thus, the nephrin-Dynll1 colocalization is used as an in vivo biomarker for dynein-mediated trafficking of nephrin. Our immunofluorescence labelling in kidney biopsy sections obtained from patients with DN revealed an enriched expression of Dynll1 along the glomerular capillaries that colocalized with nephrin (**Figure 7**). Such a pattern was also observed in patients with FSGS but is absent from the normal kidney or in MCD, a reversible podocytopathy with a benign outcome.

**Figure 7.**
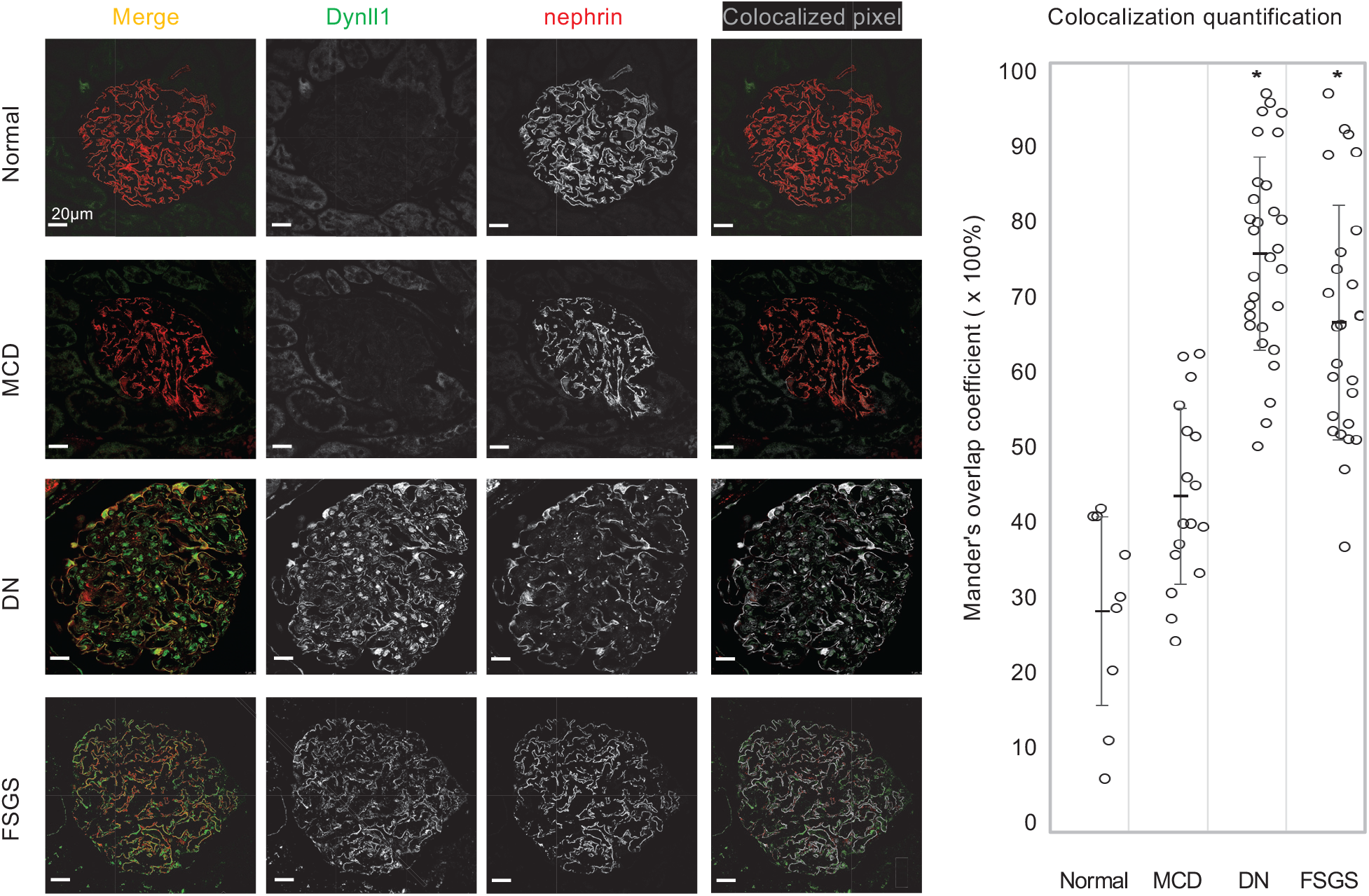
In situ biomarker for dynein mediated trafficking of nephrin in human glomerulopathies. Colocalization of Dynll1 and nephrin under conditions of increased Dynll1 expression was examined by co-immunofluorescent staining and was compared in progressive podocytopathies (FSGS and DN) vs. normal kidney and benign podocytopathy (MCD) (Scale Bars = 20µm). The amount of Dynll1 that colocalized with nephrin in non-sclerotic glomeruli was quantified using the *Colo2* plugin of Fiji software as *Manders coefficients*, and these parameters were compared among DN (n= 30 glomeruli in 10 specimens), FSGS (n= 26 glomeruli in 10 specimens), MCD (n= 17 glomeruli in 7 specimens) and normal kidney (n=9 glomeruli in 3 specimens). \**p<0*.*05* vs. normal kidney.

We further tested whether the significant Dynll1-nephrin colocalization in human DN is related to an altered sequestration of INF2. Compared to normal control kidney specimens, there is a significant colocalization of Dynll1 with INF2 along the edges of podocytes in patients with DN (**Figure 8**). Although the expression of INF2 was increased in DN, this level of upregulation is far less dramatic than that of Dynll1 (**Figure 1C&D**), suggesting inadequate sequestration of Dynll1 by INF2 in diabetic podocytopathy.

**Figure 8.**
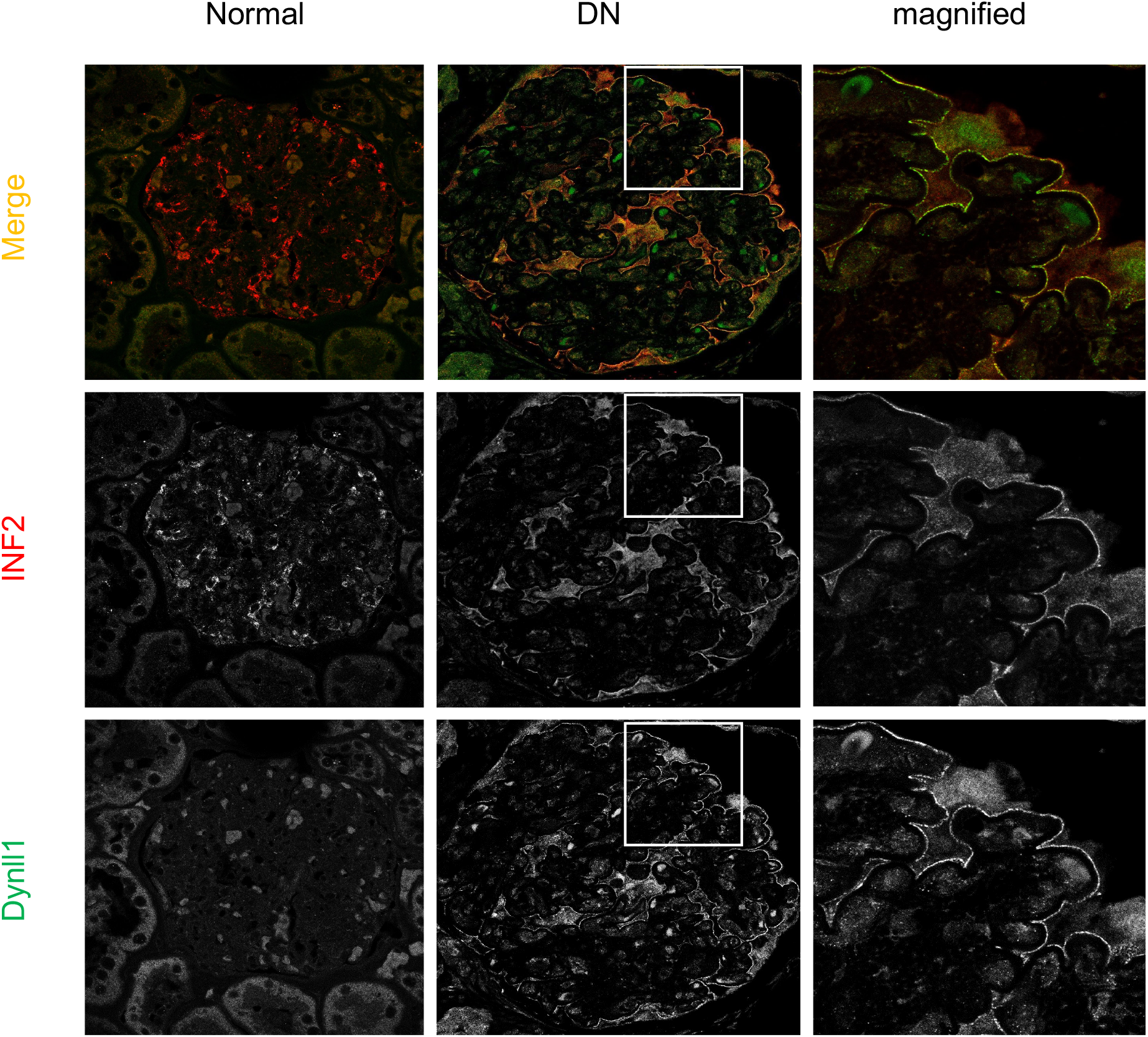
Dynll11 protein expression is increased and colocalizes with INF2 in human DN.

## Discussion

DN is a nearly ubiquitous end organ-complication of diabetes, reflective of the fact that chronic hyperglycemia is a major cause of podocyte injury. This injury results in increased GFB permeability evidenced by the development of albuminuria, followed by the progression to glomerulosclerosis ^10, 49-51^. The ability to identify the earliest pathophysiological events of DN and intervene in these before the development of diabetic kidney injury holds the greatest promise for improving disease outcome. It is well known that nephrin plays a critical role in GFB architecture. Loss of the precise cell surface membrane targeting of nephrin results in podocyte foot process effacement and SD disassembly. Increased nephrin endocytosis has been described in diabetic podocyte injury^14, 15^; however, our understanding of the molecular and cellular events driving nephrin loss are not clear. Consequently, we do not yet have a complete explanation for how the diabetic milieu changes the homeostasis of key podocyte proteins, nor a good therapeutic target to interfere with this process. Our study reveals a new mechanism that the turnover of nephrin in DN is mediated by dynein-driven, microtubule-dependent trafficking that diverts internalized nephrin towards lysosomal degradation, and thus inspires novel therapeutic approaches.

In this study, we started by analyzing multiple DN transcriptome datasets collected in *Nephroseq* kidney transcriptome databases. We discovered elevated expression of genes encoding dynein components that correlated positively with hyperglycemia, as well as with the development of albuminuria and renal insufficiency, providing *in vivo* evidence to support the potential involvement of dynein in the pathogenesis of DN, especially hyperglycemia-triggered injury. *In vitro* studies found podocytes grown in hyperglycemic conditions recapitulate a number of signaling mechanisms leading to podocyte injury in *in vivo* models of type I and tyle II diabetes, including evoking AMPK-related insulin resistance^52^ reactive oxygen species^53^, Rapamycin-Induced Endoplasmic Reticulum Stress^54^. We verified that expression of dynein components is elevated in podocytes growing in hyperglycemic conditions and in an STZ-induced type I diabetic mouse model characterized by prominent hyperglycemia; these results support the role for dynein in mediating the early pathophysiology of DN induced by hyperglycemia. Specifically, our data highlight a prominent role for *DynII1* and *DCTN1* as key responders to hyperglycemia, which can be explained by the fact that their promoters harbor binding motifs for hyperglycemia-responsive transcription factors like Specificity Protein 1 (SP1) and Krüppel-like factors^55^. Further studies are needed to clarify the hyperglycemia mediated cell signaling related to the activation of these transcription factors. *DCTN1* encodes dynactin 1, a key subunit of the dynactin complex, which is a required activator for dynein-mediated retrograde trafficking^23^; *Dynll1* encodes the light chains that play a pivotal role in maintaining the integrity of the entire dynein complex and its activity, as evidenced by the observation that dynein-mediated processes can be limited by molecular traps for Dynll1 protein.^24, 45^ Our previous research found that inadequate sequestration of Dynll1 by R218 mutant of INF2 disturbs dynein mediated trafficking of nephrin in podocytes and leads to FSGS^28^. Our immunostaining showed a significant colocalization of Dynll1 with INF2 in podocytes of diabetic patients, suggesting that the intracellular INF2 is trying to sequester the overexpressed Dynll1. However, INF2 expression does not increase to the high levels that Dynll1 does in STZ-induced DN and is likely inadequate to temper the effects of overexpressed Dynll1.

Similar to our findings, in another model of type I DN (the OVE26 mouse), *Dynll1* was found to be one of the most overexpressed genes in the kidney^56^. The expression of dynein in any type II DN animal model has not been formally reported. Our reanalysis of the Hodgin Diabetes Mouse Glom dataset revealed dynein overexpression in two type II DN models, *db/db* mice and *eNOS-/- db/db* double mutant mice, providing a strong rationale for further focused studies to test the role this mechanism in type II DN models in the future.

To test whether the hyperglycemia-induced dynein overexpression upregulates dynein activity in podocytes, we analyzed our *in vitro* nephrin trafficking model and observed increased recruitment DCTN1 and Dynll1 to the endocytosed nephrin in HG-treated podocytes, suggesting an enhanced involvement of dynein in the post-endocytic sorting of nephrin. This mechanism was also evidenced *in vivo* by the observed enrichment of Dynll1 and DCTN1 in nephrin pulldowns in STZ-induced mouse DN and reflected by increased expression of Dynll1 that colocalized with nephrin in human DN.

Furthermore, the recruitment of HDAC6 to endocytosed nephrin in HG-treated cells indicates the sorting of these nephrin to lysosomal degradation pathways, since HDAC6 functions as an adaptor that connects dynein with ubiquitinated cargo proteins and diverts them to lysosomes^42, 43^. This was supported by the increased flux of endocytosed nephrin to Rab7-positive late endosomes/lysosomes in HG-treated cells, as well as by the fact that the reduced nephrin protein could be salvaged by inhibiting lysosomal proteases using Leupeptin. Meanwhile, under hyperglycemic conditions, we found decreased sorting of nephrin to Rab11-positive recycling endosomes and impaired surface recycling of nephrin, which proved to be a dynein-dependent change. The fact that these changes could be rescued by inhibiting dynein ATPase with Ciliobrevin D or by knocking down *Dynll1* demonstrated the key role of dynein, especially Dynll1, in hyperglycemia-induced mistrafficking of nephrin. These findings support the hypothesis that diabetic podocytopathy occurs when hyperglycemia activates the expression of dynein, which in turn facilitates the depletion of nephrin by causing its mistrafficking away from recycling and into degradation pathways through adaptors like HDAC6 ^42^. Since nephrin is an integral membrane protein, these findings suggest that dynein-mediated post-endocytic sorting of nephrin results in its degradation, primarily by lysosomes via sorting through multi-vesicular bodies. Although our data also showed the stabilization of endocytosed nephrin by Bortezomib, a proteasome inhibitor, it is not possible to conclude that the proteosome also plays a significant and direct role in the post-endocytic degradation of nephrin, since there is evidence suggesting that proteasome inhibitors can sequestrate available ubiquitin and subsequently limit the ubiquitin-dependent degradation in lysosomes, too.

In our *in vivo* model of STZ-induced DN, we observed increased ubiquitinated nephrin and, reduced surface and total nephrin but without reduced transcription of *NPHS1*. These changes corresponded to the overexpression of dynein components that were recruited to nephrin. Moreover, we provided *in vivo* evidence of dynein-mediated trafficking of nephrin in patients with DN, similar to FSGS, but not in MCD, indicating dynein may mediate certain processes that distinguish deteriorating podocytopathies from reversible ones. Meyer-Schwesinger *et al*’s work revealed an upregulated ubiquitin proteolysis system underlying the irreversible loss of podocyte proteins in DN and FSGS, compared to MCD^57^. The unique role of dynein in bridging the initial endocytosis of podocyte proteins to the end process of proteolysis provides a potential explanation for the progressive nature of podocyte injury in diabetes.

Taken together, our work defines a mechanistic role for dynein-mediated, post-endocytic sorting of nephrin in the pathogenesis of DN. By diverting nephrin from surface recycling to degradation, dynein-mediated sorting provides a viable explanation for the loss of nephrin that compromises the integrity of the SDs in DN. Recognizing the role of dynein-driven mistrafficking and degradation of nephrin opens the door for the development of new therapies for human DN. Specifically, the therapeutic opportunities include modifiers targeting specific proteins or relevant cellular events, including dynein activity, ubiquitin ligases or deubiquitinating enzymes, and HDAC6 activity, as well as therapeutics for the modulation of microtubule polymerization and the signal transduction pathway(s) that connects hyperglycemia with dynein expression/activation.

## Data Statement

The transcriptome data were derived from the Hodgin Diabetes Mouse Glom dataset, Woroniecka Human Diabetic Kidney Disease dataset and the European renal cDNA bank nephrotic syndrome dataset collected by the Nephroseq data-mining platform (www.nephroseq.org, University of Michigan, Ann Arbor, MI). The other data supporting the findings can be requested from the corresponding author.

## Acknowledgments

The authors would like to acknowledge use of the University of Iowa Central Microscopy Research Facility (CMRF), a core resource supported by the University of Iowa Vice President for Research, and the Carver College of Medicine. We thank the advisory committee of CMRF for reviewing the microscopy protocols. We thank the Department of Pathology and Pollak Laboratory, Beth-Israel Deaconess Medical Center (BIDMC) for sharing normal human kidney sections. We thank Mr. Aniket Gad at the BIDMC confocal core facility for his help and technical support on confocal microscopy. We thank Mr. Andrew Costello and Ms. Mariah R Leidinger at the Pathology Department, the University of Iowa Hospitals and Clinics for their help in the experiments. We thank Dr. Michael R Rebagliati for the scientific editing of this manuscript.

## Author contributions

HS and JW were responsible for the design and execution of experiments, statistical analysis, and drafting of the manuscript. CA was responsible for the confocal microscopy and contributed to the design, protocol development and quantitative imaging data analysis. RP contributed to the research design, data review and interpretation, as well as writing of the manuscript. JM contributed to the writing and critical editing of the manuscript. CN contributed to the design of the human pathology study and result interpretation, as well as the writing of the manuscript. All authors approved the final version of the manuscript.

## Figures legends

**Supplementary Figure 1. Correlation of dynein expression with diabetic nephropathy phenotypes**.

Transcription analyses of the Hodgin Diabetes Mouse Glom dataset showed that the expression levels of dynein components correlated with microalbuminuria (albumin to creatinine ratio, a) and fasting glucose levels (b) in various diabetic mouse models (*eNOS*^*-/-*^ *db/db* mice, streptozotocin-induced diabetic mice, and *db/db* mice). c. Transcription analyses of the Woroniecka Human Diabetic Kidney Disease dataset and ERCB Chronic Kidney Disease dataset revealed that the expression of dynein components correlated inversely with the glomerular filtration rate (GFR) in human diabetic nephropathy.

