## Supplementary Figure 1 for "Dynein-mediated trafficking and degradation of nephrin in diabetic podocytopathy"

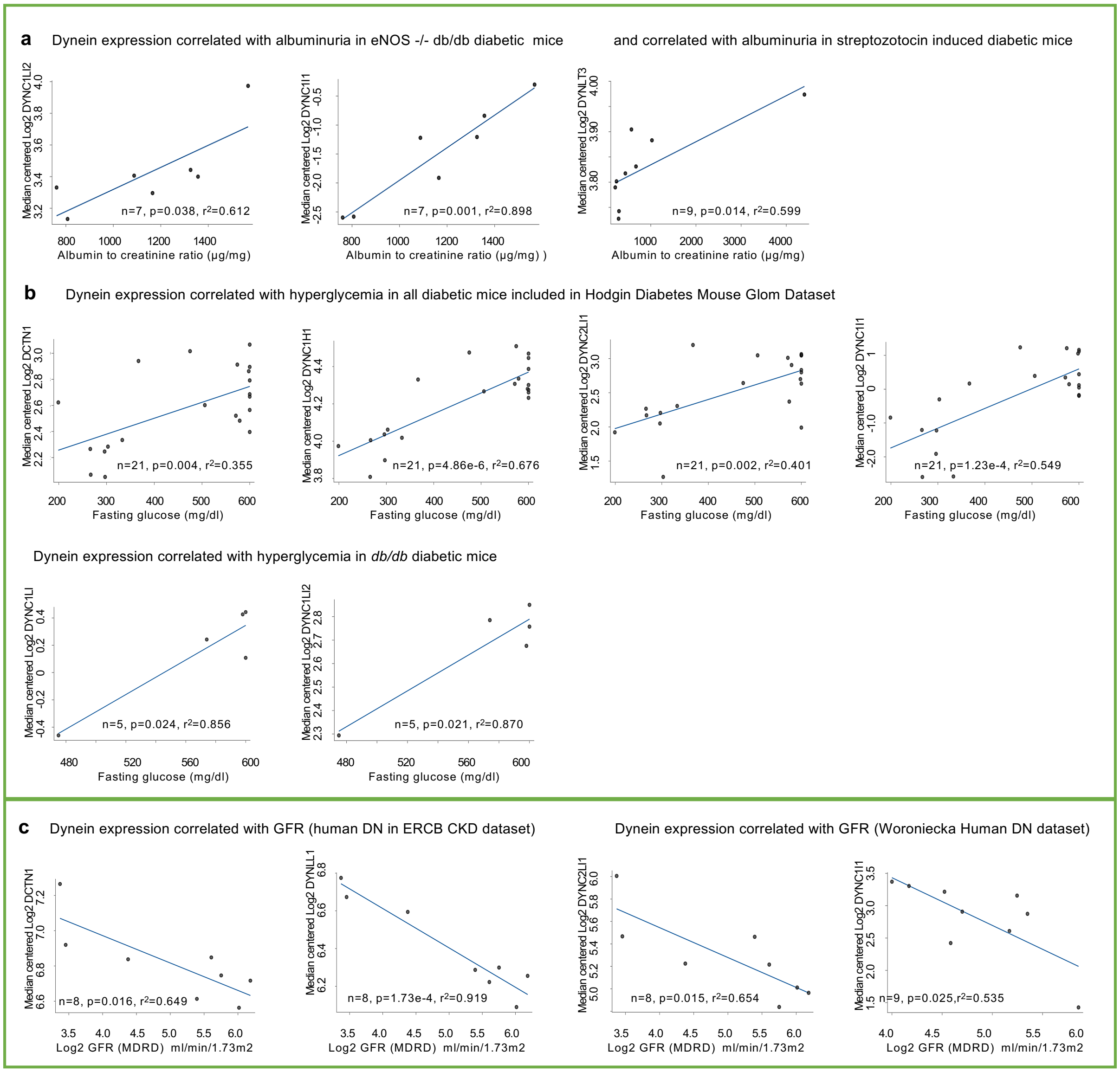


**Supplementary Figure 1*.* Correlation of dynein expression with diabetic nephropathy phenotypes.** Transcription analyses of the Hodgin Diabetes Mouse Glom dataset showed that the expression levels of dynein components correlated with microalbuminuria (albumin to creatinine ratio, a) and fasting glucose levels (b) in various diabetic mouse models (*eNOS*$^{-/-}$ *db/db* mice, streptozotocin-induced diabetic mice, and *db/db* mice). c. Transcription analyses of the Woroniecka Human Diabetic Kidney Disease dataset and ERCB Chronic Kidney Disease dataset revealed that the expression of dynein components correlated inversely with the glomerular filtration rate (GFR) in human diabetic nephropathy.
